# CRISPR-mediated biocontainment

**DOI:** 10.1101/2020.02.03.922146

**Authors:** Oscar Castanon, Cory J. Smith, Parastoo Khoshakhlagh, Raphael Ferreira, Marc Güell, Khaled Said, Ramazan Yildiz, Matthew Dysart, Stan Wang, David Thompson, Hannu Myllykallio, George M. Church

**Affiliations:** Department of Genetics, Harvard Medical School, Boston, Massachusetts, USA; Wyss Institute for Biologically Inspired Engineering, Boston, Massachusetts, USA; LOB, Ecole Polytechnique, CNRS, INSERM, Institut Polytechnique de Paris, 91128 Palaiseau, France; Pompeu Fabra University, Barcelona Biomedical Research Park, 08003-Barcelona (SPAIN)

## Abstract

We have exploited the repetitive nature of transposable elements of the human genome to generate synthetic circuits. Transposable elements such as LINE-1 and Alu have successfully replicated in mammalian genomes throughout evolution to reach a copy number ranging from thousands to more than a million. Targeting these repetitive elements with programmable DNA nucleases such as CRISPR-Cas9 rapidly induce extremely high levels of cell death. We use this genotoxic feature to build synthetic biocontainment circuits: CRISPR defense system (CRISPR-DS) capable of preventing CRISPR genome editing, and we introduce the proof-of-concept of CRISPR Safety-Switch, an inducible, stringent and non-leaky kill-switch capable of clearing out cell lines resistant to DNA breaks.

## Introduction

Double strand breaks (DSB) are toxic in prokaryotic and eukaryotic cells. Generating a single DSB with programmable nucleases in prokaryotic genome has been shown to trigger cell death^1^ and have been efficiently used as a selection system in synthetic systems^2^ or as potential antimicrobial strategies^3^. In contrast, mammalian systems have evolved more efficient DSB DNA repair mechanisms such as non-homologous end joining (NHEJ) and homology directed repair (HDR) to heal discrete injuries to the genome^4,5^. When excessive damage has been done to the DNA, eukaryotic cells trigger apoptotic pathways^6^ to prevent the mutations from disrupting normal cell functions or spreading to the next generations, potentially triggering disease. Several studies have reported the deleterious effects of targeting sites in moderately repetitive regions (4-62 copies per cell)^7,8,9^ using programmable nucleases such as CRISPR/Cas9. While Kuscu et al. reports elevated apoptosis at ~12 copies per cell, higher amount of double-stranded breaks has been shown to be well tolerated in at least some cells as demonstrated by the isolation of multiple independent clones with the knock-out of 62 Porcine Endogenous Retroviruses (PERV) elements resulting in viable cells and ultimately the birth of healthy pigs without PERV expression or transmission. To achieve high Cas9 genotoxicity and create synthetic biocontainment tools, we explored the targeting of transposable elements naturally found within eukaryotic genomes. Transposable elements are widespread repetitive sequences in many organisms’ genomes that can duplicate and/or move to a new locus, either autonomously or dependent on another mobile element^10^. The initial publication of the human genome^11,12^showed that up to two-thirds of the nuclear DNA was repetitive^3^ and is primarily transposon derived. The Long-Interspersed Elements-1 (LINE-1) and the Alu elements are the most widespread transposable elements which respectively constitute about 17% and 10% of our genome^10^.

While attempting large-scale genome editing at these transposable elements, we have previously shown that targeting LINE-1 sequences in the genome using the original CRISPR-Cas9 system in HEK 293T, did not yield the survival of any edited clones^14^. Our hypothesis was that Cas9 would trigger a massive number of DSBs at these targets, overwhelming the cell’s ability to repair this damage. Therefore, we further tested single guide-RNAs (sgRNAs) targeting LINE-1 and Alu, respectively displaying repeat numbers of about 10^4^ copies^15^ to 10^6^ copies^16^.

In this study, we aim at generating a system that can stringently eliminate cells on demand and which could be implemented in a variety of useful eukaryotic synthetic circuits. As genome editing forces its way into society with a potential for harmful applications, the research community must reflect upon its impact in our lives and develop potential countermeasures. Here, we designed CRISPR defense system (CRISPR-DS) to prevent CRISPR-Cas9 editing as a novel way to shield genomes from unwanted genetic modification. We tested our circuit against systems using sgRNAs targeting the essential genes *POLE2* (sgRNA-*POLE2*), a subunit of the DNA polymerase, and *GTFIIB* (sgRNA-*GTFIIB*), the General Transcription factor IIB and anti-CRISPR proteins^17^, bacteriophages molecules that have naturally evolved to inhibit Cas9 nucleases. CRISPR-DS showed to be more stringent than such systems but they could potentially be used in a complementary manner.

In addition, although the field of cell therapy advances and continues to show great promise to cure a wide variety of diseases^18^, several risks remain, notably, the potential of engineered cells to become oncogenic or to trigger cytokine release syndrome in the context of CAR-T therapies. In order to mitigate these risks, we generated the proof-of-concept of an inducible, stringent and non-leaky safety switch as a new approach to selectively eliminate transplanted therapeutic cells in case of adverse effects.

## Methods

### Cas9, sgRNA and anti-CRISPR plasmids used for genome editing

Expression vector encoding humanized pCas9_GFP protein was obtained from Addgene.org (Plasmid #44719). The traditional route of using online sgRNA design tools does not work for repetitive elements as they are designed to avoid these sequences for concerns of deleterious off-target effects. sgRNAs were therefore manually designed on the most conserved segments of the consensus sequences (Suppl. Table 2). The consensus sequence for Alu was obtained from the repeatmasker database (http://www.repeatmasker.org/species/hg.html) and the consensus for LINE-1 was based on the 146 intact full length elements from the L1base^15^. The sgRNAs used in this study were synthesized and cloned as previously described^19^, briefly two 24mer oligos with sticky ends compatible for ligation were synthesized from IDT for cloning into the pSB700_mCherry plasmid (Addgene Plasmid #64046) after cutting with the BsmbI restriction enzyme. After sequence confirmation using the humanU6 primer, plasmids were prepared using the Qiagen Plasmid Plus Midi Kit (Cat # 12943). Expression vectors encoding Anti-CRISPR proteins were obtained from Addgene: AcrIIA2 (pJH373 plasmid, ID# 86840) and AcrIIA4 (pJH376 plasmid, ID# 86840).

### Synthesis and genomic integration of the Cas9 and sgRNAs into HEK 293T cells

sgRNAs targeting Alu, LINE-1, and a non-human control were amplified from the pSB700 plasmid and cloned into PB-TRE-dCas9_VPR (Addgene #63800) using the following primers: U6-NheI-F, and sgRNA-*Bam*HI-R (Suppl. Table 3). dCas9_VPR was removed during the cloning process using the restriction enzymes *Nhe*I and *Bam*HI to integrate the U6-gRNA construct into the PiggyBac transposon sequences. Colonies were Sanger sequence verified and prepped using the Qiagen plasmid plus midi kit. HEK 293T cells were then lipofected with PB-gRNAs (Alu, AluYa5, LINE-1, and non-human) and PB-transposase. Cells were selected with puromycin (1μg/ml) beginning at day two until day nine. Populations of puromycin resistant cells were used for the initial Cas9 genome editing trials. Individual cells from the puromycin resistance population were grown after single-cell sorting into 96-well plates and isolated for further testing in Cas9 genome editing experiments.

### Propidium Iodide and Annexin V staining and FACS analysis

Cells were dissociated with TrypLE, diluted in an equal volume of PBS then centrifuged at ~300g for 5 minutes at room temperature. We resuspended samples into 500μl PBS and half of the cells were pelleted for later gDNA analysis. The remainder was centrifuged and resuspended into 100μl of Annexin V Binding Buffer (ref #V13246) diluted into ultrapure water at a 1:5 ratio. Subsequently, we added 5μl of Alexa 647 Annexin V dye (ref #A23204) and incubated samples in the dark for 15 minutes. We then added 100μl of Annexin V Binding Buffer and added 4μl of Propidium Iodide (ref #P3566) diluted into the Annexin V Binding Buffer at a 1:10 ratio. Samples were incubated in the dark for another 15 minutes. Cells were washed with 500μl of Annexin V Binding Buffer and centrifuged again to be finally resuspended into 400μl of Annexin V Binding Buffer. All samples were filtered using a cell strainer and were run on the LSR II using a 70-μm nozzle. Subsequent analysis was conducted using FlowJo V10 software.

### Antibody staining and fluorescent microscopy

Cells were fixed with 4% formaldehyde for 10 minutes at room temperature and blocked with PBS containing 10% normal donkey serum, 0.3 M glycine, 1% BSA and 0.1% tween for 2h at room temperature. Staining of the treated cells with Anti-γH2AX antibody (10μg/ml) was performed overnight at 4°C in PBS containing 1% BSA and 0.1% tween. The cells were washed three times (5-minute intervals) with PBS followed by secondary staining. Cells were imaged using a Zeiss AxioObserver.Z1 microscope equipped with a Plan-Apochromat 20×/0.8 objective, an EM-CCD digital camera system (Hamamatsu) and a four-channel LED light source (Colibri), and Zeiss TIRF/ LSM 710 confocal (ZeissTIRF-confocal), 63×.

### Transfection of HEK 293T

HEK293T cells were cultured in Dulbecco’s modified Eagle’s medium supplemented with 10% FBS (Gibco) at 37 °C with 5% CO_2_ incubation. Transfection was conducted using Lipofectamine 2000 (Cat# 11668027) using the suppliers recommended protocol. 1μg SpCas9_GFP, 1μg sgRNA-JAK2, and 1μg sgRNA-test were used per 80000 cells in a 12-well plate. Cell pellets were collected three days after transfection for genomic DNA extraction and sequencing analysis.

### Preparation of HEK 293T samples for Insertions and Deletions (indels) analysis

Following the genomic DNA extraction of HEK293T samples using DNeasy Blood & Tissue Kit from Qiagen (Cat# 69506) according to the supplier’s protocol, we amplified 586 bp of the JAK2 locus using the Sanger-JAK2-F and Sanger-JAK2-R primers (Suppl. Table 3). Amplicons were obtained after PCR amplification using Kapa HiFi HotStart Readymix kit from Kapa Biosystems (Cat# KK2602) according to the supplier’s protocol. PCR products were then run on E-Gels EX 2% Agarose (Cat# G402002) from Invitrogen and amplicons of about 586 bp were extracted using the Qiagen QIAquick Gel Extraction Kit (Cat# 28706). Gel extracted PCR products were then submitted to Genewiz for Sanger DNA sequencing using the P3R primer.

### Insertions and deletions (indels) analysis

Indels analysis of all samples from Fig. 3a was executed using TIDE web tool^20^. The experiment was performed with 3 replicates as described in “Preparation of HEK293T samples for Insertions and Deletions (indels) analysis”. Sequencing trace files provided through Genewiz services were then analyzed using TIDE web tool that assesses genome editing by CRISPR/Cas9 using a decomposition algorithm that identifies and quantifies insertions and deletions in the expected editing site. The following advanced settings were used: Alignment window: left boundary = 100; Decomposition window = 268 bp to 350 bp; indels size range = 0 to 10 bp; P-value threshold = 0.001. For each sample, the Control Sample Chromatogram file uploaded comes from a HEK 293T sample transfected only with sgRNA-control. Data was plotted using excel displaying the mean of three biological replicates with the error bars representing the standard error. Statistical analysis was conducted using the Student’s t-test.

### Illumina MiSeq library preparation and sequencing

Library preparation was conducted as previously described^21^. Briefly, genomic DNA was amplified using locus-specific primers attached to part of the Illumina adapter sequence (Suppl. Table 3). A second round of PCR included the index sequence and the full Illumina adapter. Libraries were purified using gel extraction (Qiagen #28706), quantified using the NanoDrop and pooled together for deep sequencing on the MiSeq using 150 paired end (PE) reads.

### NGS data analysis

The activity of Cas9 was measured by the number of reads containing insertions or deletions around the sgRNA target site. FastQC was initially used to confirm sequence quality, length and diversity. CRISPR RGEN tools^22^ was used to quantify indel disruption at the targeted site by submitting the fastq file, reference, and sgRNA sequence. Data was plotted using excel displaying the mean of three biological replicates with the error bars representing the standard error. Statistical analyses were conducted using the Student’s t-test.

### Bioinformatic analysis

Alignment analysis of the sgRNA sequences to all human chromosomes was performed using the R library Biostrings v2.40.2 as previously described^23^. sgRNAs were aligned to hg38 reference genome, downloaded from Ensembl website (ftp://ftp.ensembl.org/pub/release-95/fasta/homi_sapiens/dna/). PAM sequences were retrieved by taking three nucleotides adjacent to the different alignments. Distribution of the different hits and PAM sequences was plotted using the R library ggplot2 3.3.0.

## Results

### Design and identification of lethal sgRNAs targeting transposable elements of the genome

We previously designed repetitive sgRNAs to target about 26000 human LINE-1 (sgRNA-LINE-1) and about 161000 Alu (sgRNA-Alu) elements^14^ based on their consensus sequences to generate numerous double-strand DNA breaks and initiate apoptosis (or otherwise render the cell non-viable). These lethal sgRNAs may be expressed in a cell prophylactically, being benign to the cell until it encounters the Cas9 nuclease (Fig. 1a, Fig. S1).

**Figure 1.**
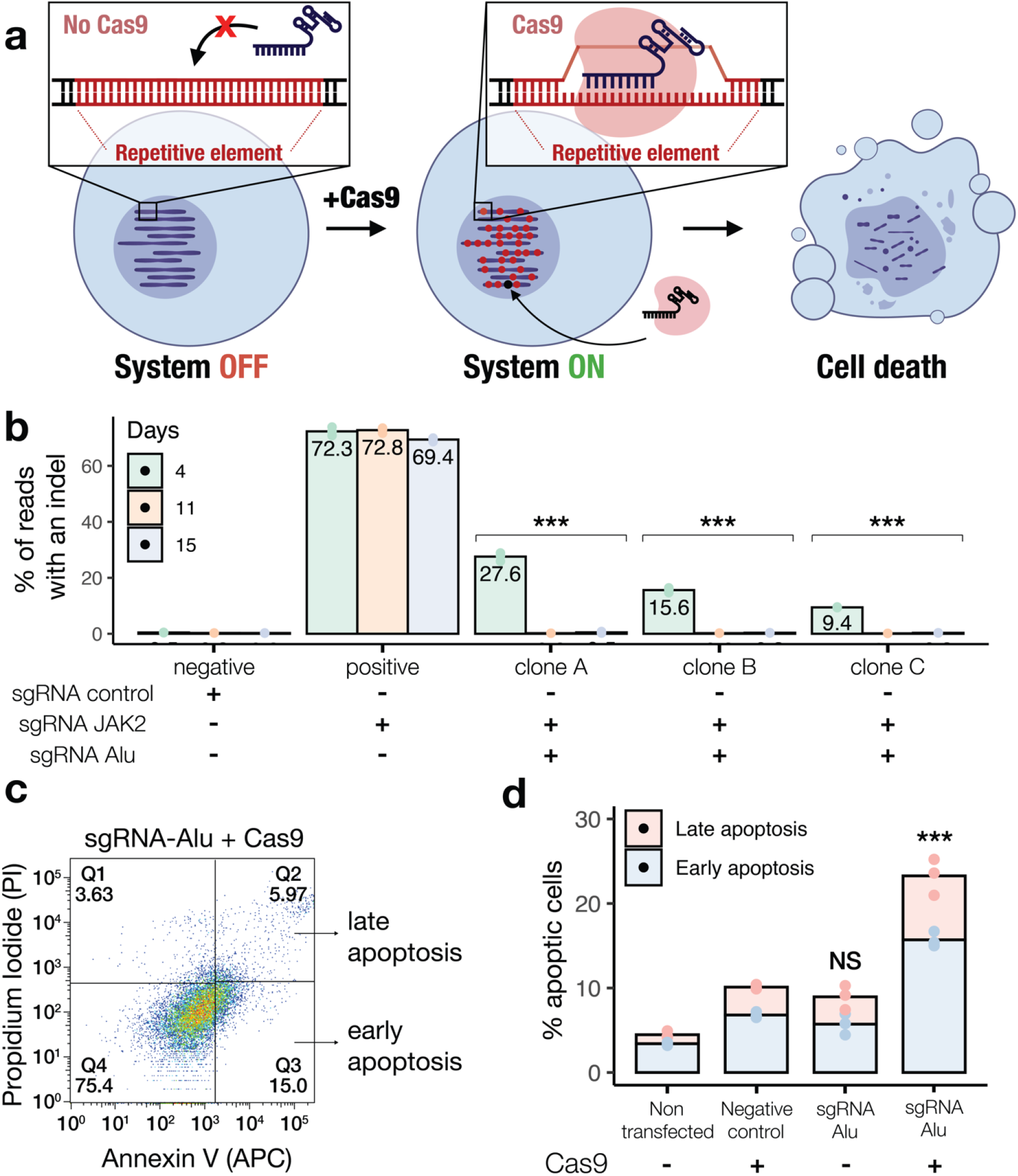
Principle of the CRISPR-Defense System (CRISPR-DS): A genomically integrated sgRNA targeting transposable elements prevents the generation of cell populations with unintended. (**a**) Schematic of typical CRISPR/Cas editing in the presence of Cas9 sensor before transfection (system OFF) and after transfection (system ON). Cas9 pairs with the sgRNA and disrupts its target site creating indels. The chromosomal sgRNA targeting repetitive elements is benign in the absence of Cas9 but acts as a surveillance system for incoming Cas9, resulting in cell death. (**b**) Prevention of DNA modification at the JAK2 locus. The graph represents the mean of three biological replicates for indel mutation rate at the JAK2 locus at days 4, 11, and 15 after transfection in light green, orange, and blue respectively. (**c**) Apoptosis assay FACS plot of the sample treated with sgRNA-Alu. The plot separates early apoptotic HEK 293T cells (Annexin V+ propidium iodide-) from late apoptotic HEK 293T cells (Annexin V+ propidium iodide+). (**d**) Early (light blue) and late apoptotic cells (light orange) percentages of the samples with (+) or without (−) Cas9 three 3 days after transfection. In all histograms, error bars represent standard error, n=3. Student’s t-test was performed and marked NS, not significant (P > 0.05); *P < 0.05; **P < 0.01; ***P < 0.001 as compared to the positive control in (b) and to the negative control in (d).

In theory, such a synthetic system should decrease genome editing at a known high-efficiency locus – such as *JAK2* – since every cell co-transfected with a repetitive sgRNA should trigger apoptosis, ultimately clearing the cells harboring any edit at the *JAK2* locus. To test this, we transiently transfected sgRNA-Alu or sgRNA-LINE-1, along with *Streptococcus pyogenes* Cas9 (SpCas9) and a sgRNA targeting *JAK2* – our test gene – in Human Embryonic Kidney 293T (HEK 293T) cells. We confirmed with Next Generation Sequencing (NGS) that sgRNA-Alu and sgRNA-LINE-1 decreased *JAK2* insertions and deletions (indels) by about 3.6-fold two days after transfection, compared to our positive control (Fig. S2). With similar efficiencies between our two repetitive sgRNAs, we decided to proceed with sgRNA-Alu in designing our synthetic circuits as it targets significantly (a 10-fold difference) more sequences in the genome than sgRNA-LINE-1.

### CRISPR-Defense System: A genomically integrated repetitive sgRNA prevents the formation of populations harboring DNA edits

For our next phase of testing, we integrated constitutively expressed repetitive element-targeting sgRNAs into the genome using PiggyBac (PB) transposition in HEK 293T. We assayed DNA editing efficiency by transfection with SpCas9 and a sgRNA targeting our test gene *JAK2* (sgRNA-*JAK2*). Clonal cell populations stably expressing repetitive element targeting sgRNAs were isolated and exposed to this genome editing challenge. DNA was subsequently analyzed with NGS at several different time points to quantify the presence of edits at the *JAK2* locus (Fig. 1b). With exposure to SpCas9, any successfully *JAK2*-modified cell will also be cut at the high-copy repetitive element, resulting in rapid and near complete cell clearance. In HEK 293T cells, while non-homologous end joining (NHEJ) levels of up to 72% were observed in control samples, the three sgRNA-Alu expressing clonal cell populations A, B and C, displayed 27.6%, 16% and 9.4% indels by day 4 and background levels of 0.10%, 0.09% and 0.04% by day 15, respectively, representing a ≥99.9% reduction of DNA editing in clones expressing sgRNA-Alu as compared to control cells stably expressing a sgRNA targeting no sequences in the human genome (sgRNA-control). These results show that, as expected, the biocontainment system is effective at preventing cell populations from being genetically altered by clearing out cells in which SpCas9 is expressed.

### CRISPR-Cas9 targeting high-copy number loci rapidly causes DNA damage

Evidence that cells expressing CRISPR-DS in these experiments are removed following Cas9 expression is supported by the decrease over time of fractions of cells exhibiting *JAK2* mutations; however, these data do not identify the mechanism of cell death. We hypothesized that the cells containing our synthetic system undergo apoptosis triggered by the massive DNA damage caused by the expression of SpCas9^7^. We tested this hypothesis by undertaking standard cell death and apoptosis assays followed by flow cytometry analysis, and immunostaining cell samples for γH2AX, a known marker of DSBs. In HEK 293T cells stably expressing sgRNA-Alu, expression of SpCas9 significantly increased the percentage of early apoptotic and late apoptotic cell populations, as measured by Annexin V and propidium iodide, exceeding by about 2.3-fold the apoptosis triggered in HEK 293T control cells stably expressing the negative control sgRNA (Fig. 1c, 1d). On the contrary, when HEK 293T cell lines expressing sgRNA-Alu did not receive SpCas9, they displayed similar cell death levels to cells expressing sgRNA-control (Fig. 1c, 1d) showing that sgRNA-Alu did not display any abnormal toxicity on its own. With respect to the γH2AX staining associated with DSB induced DNA damage, we observed a clear increase in γH2AX foci along with the abnormal formation of fused cells in the sgRNA-Alu expressing cells that was not observed in the non-human targeting control (Fig. 2a, 2b, 2c). These results support the hypothesis that repetitive sgRNAs induce apoptotic death from massive simultaneous double-stranded DNA cleavage while remaining non-toxic in the absence of a foreign Cas9-based DNA editor.

**Figure 2.**
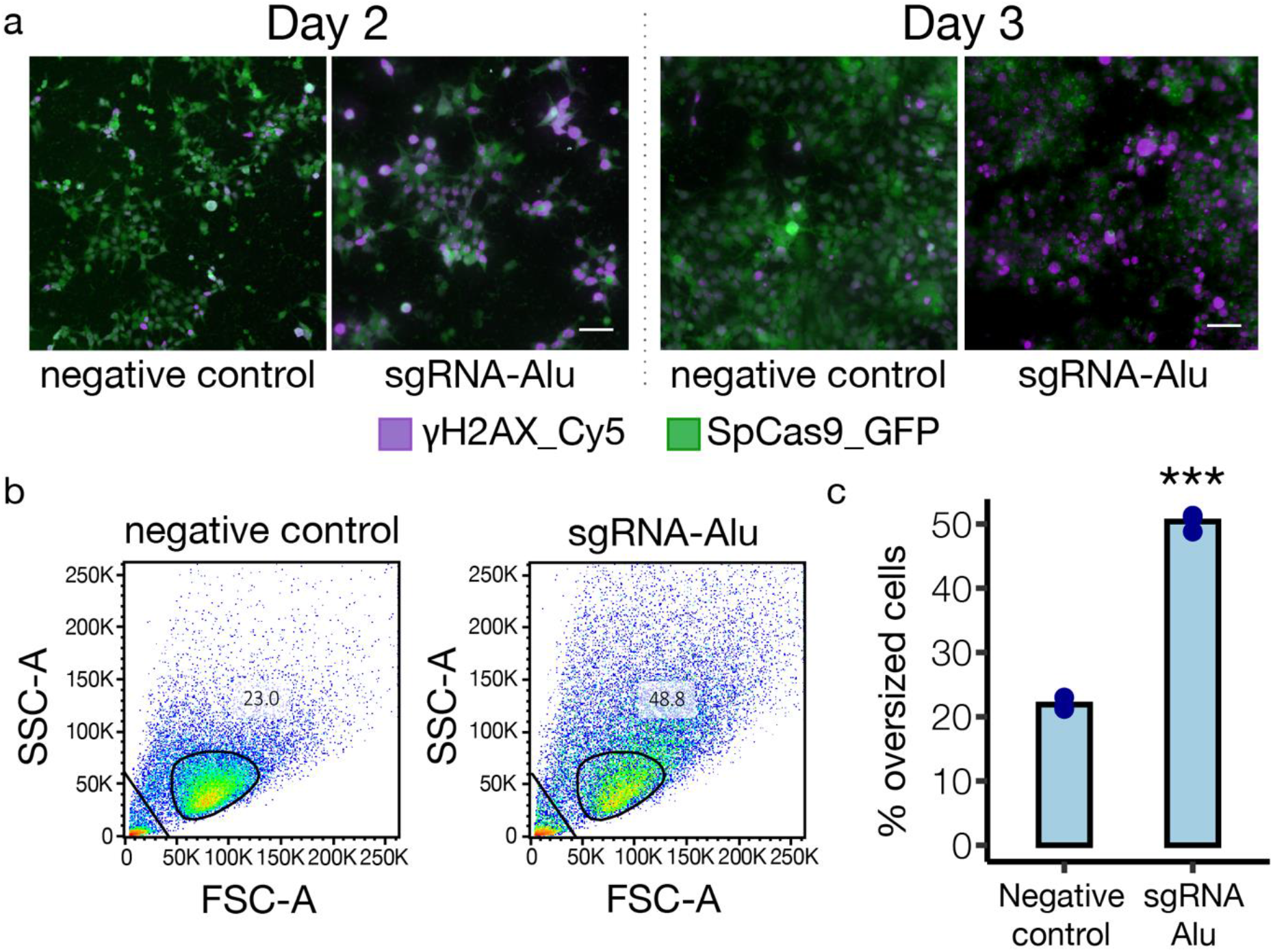
CRISPR-Cas9 targeting high-copy number loci causes DNA damage. (**a**) 20x magnification microscope images of *γ*H2AX immunostained cells that were transfected with SpCas9 and either sgRNA-Alu or sgRNA-control (negative control) 2 or 3 days after transfection. Transfected cells appear green due to the GFP marked SpCas9 (SpCas9_GFP) and *γ*H2AX foci appear purple as antibodies are stained with a Cy5 fluorophore (*γ*H2AX _Cy5). Scale bar = 50 μm. (**b**) FACS plot showing “oversized” cells using their forward scatter (FSC) on the x-axis and their side scatter (SSC) on the y-axis. Cells were analyzed three days after being transfected with sgRNA-Alu or sgRNA-control. Cells are considered “oversized” when they fall outside the normal range shown by the circular gate. Debris are excluded from the analysis (triangular gate). (**c**) Percentage of oversized cells three days after being transfected with sgRNA-Alu or sgRNA-control as measured by FACS. Error bars represent standard error, n=3. Student’s t-test was performed and marked NS, not significant (P > 0.05); *P < 0.05; **P < 0.01; ***P < 0.001 throughout the figure.

### Comparing CRISPR-DS to circuits targeting essential genes or using anti-CRISPR proteins

We next sought to compare the efficiency of CRISPR-DS in preventing DNA edits against other approaches such as using sgRNAs targeting known essential genes (e.g. DNA polymerase subunits) and using anti-CRISPR proteins (acrIIA2 and acrIIA4) to inhibit Cas9 activity in human cells. To test these systems, we transiently transfected HEK 293T cells with SpCas9 and *JAK2*-targeting sgRNA, along with either the repetitive element targeting guides sgRNA-Alu (Fig. 3a) and sgRNA-LINE-1 (Fig. S2); the essential genes *POLE2* (Fig. 3a) and *GTFIIB* (Fig. S2); the anti-CRISPR proteins AcrIIA4 (Fig. 3a) and AcrIIA2 (Fig. S2). Cells transfected with SpCas9 and our sgRNA-control alone constituted our negative control and when transfected in addition to sgRNA-*JAK2*, our positive control. The samples transfected with sgRNA-Alu showed a significant drop in the percentage of indels at *JAK2,* from 34.4% in the positive control to a background level of 0.9% by day nine after transfection (Fig. 3a). The anti-CRISPR protein AcrIIA4 was also able to decrease *JAK2* edits down to 2.6%, which was above background levels and did not display any toxicity as compared to the negative control when assayed for cell death (Fig. 3b and 3c). On the contrary, AcrIIA2 did not have any effect in our hands and could not prevent DNA edits when compared to the positive control (Fig. S2). The sgRNA-*POLE2* decreased genome editing down to 8.6% and did not show increased cell death as compared to sgRNA-control three days after transfection (Fig. 3b and 3c). Overall, CRISPR-DS using repetitive element targeting showed the highest efficiency and resulted in frequencies of edits indistinguishable from background levels. Even though essential-gene-targeting sgRNAs and anti-CRISPR protein AcrIIA4 did not lower the frequency of edits in human cells down to background levels of detection, such strategies in combination with repetitive elements targeting sgRNAs could be used to generate a multi-layered security system to safeguard the genome from DNA edits.

**Figure 3.**
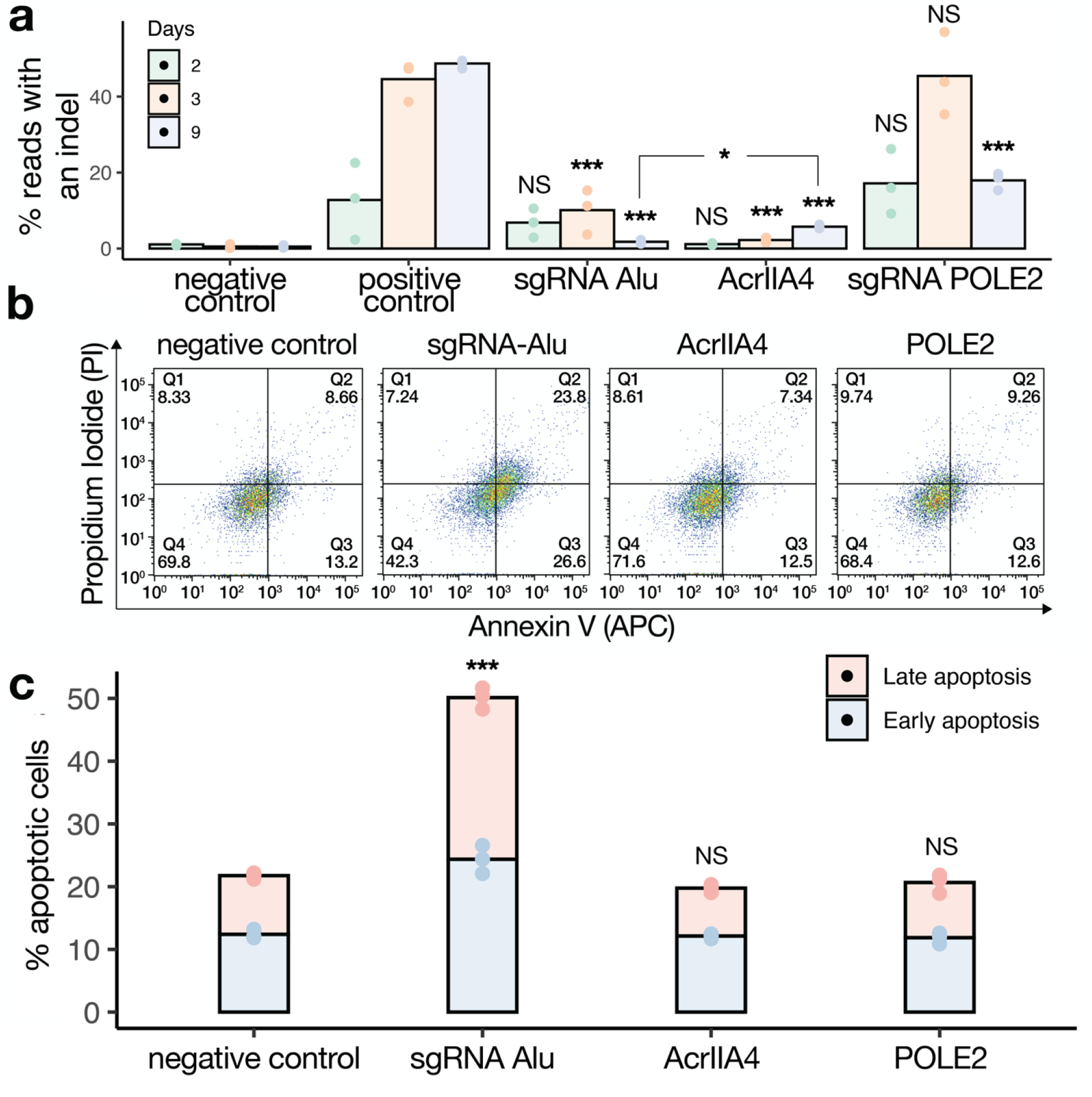
CRISPR-DS compared to systems targeting essential genes or using anti-CRISPR proteins. (**a**) Prevention of DNA edits in HEK 293T cells at the JAK2 locus by sgRNA-Alu, anti-CRISPR protein AcrIIa4, and sgRNA targeting essential gene POLE2. The percentage of reads with an indel is plotted on the y-axis and represents the mean of three biological replicates. (**b**) Three days after transfection, apoptosis was measured by FACS using Annexin V and propidium iodide staining in HEK 293T cells transiently expressing Cas9 and sgRNA-Alu, AcrIIa4, or sgRNA-POLE2. The FACS plots showing Annexin V on the x-axis and propidium iodide on the y-axis in HEK 293T are displayed for each sample. (**c**) Percentage of apoptotic cells is plotted on the y-axis with early apoptosis in light blue as measured by Annexin V+ propidium iodide-, and late apoptosis in light orange as measured by Annexin V+ Propidium iodide+ FACS populations. In all histograms, error bars represent standard error, n=3. Student t-tests were performed and marked NS, not significant (P > 0.05); *P < 0.05; **P < 0.01; ***P < 0.001 as compared to the positive control in (a) and to the negative control in (c).

### Towards the development of CRISPR Safety Switch

Leveraging the efficiency of the previously described synthetic system to eliminate human cells, we sought to build a conditional circuit to activate the cell-killing mechanism on demand. To do so, using the PiggyBac (PB) transposition system, we permanently integrated a doxycycline (DOX) inducible Cas9 endonuclease plasmid containing a hygromycin resistance cassette into the HEK 293T clonal cell line stably expressing the gRNA-Alu that showed the best cell-clearance efficiency (clone C, Fig. 1b). We expect the addition of DOX into the cell culture media of the resulting cell line to trigger Cas9 expression, enabling the targeting of the Alu elements and therefore activating the elimination of the cells containing this safety switch.

Following the described PB integration and the subsequent hygromycin selection, we obtained a cell population stably expressing both the inducible Cas9 and gRNA-Alu. Single cells of this heterogeneous population were then sorted using flow cytometry which resulted in the growth of 24 clones. Each clonal population was duplicated and treated either with or without DOX for 10 days. We selected, expanded and further analyzed the 2 clones (A’ and B’) which displayed the most cell elimination under the microscope (Fig. 4a).

**Figure 4.**
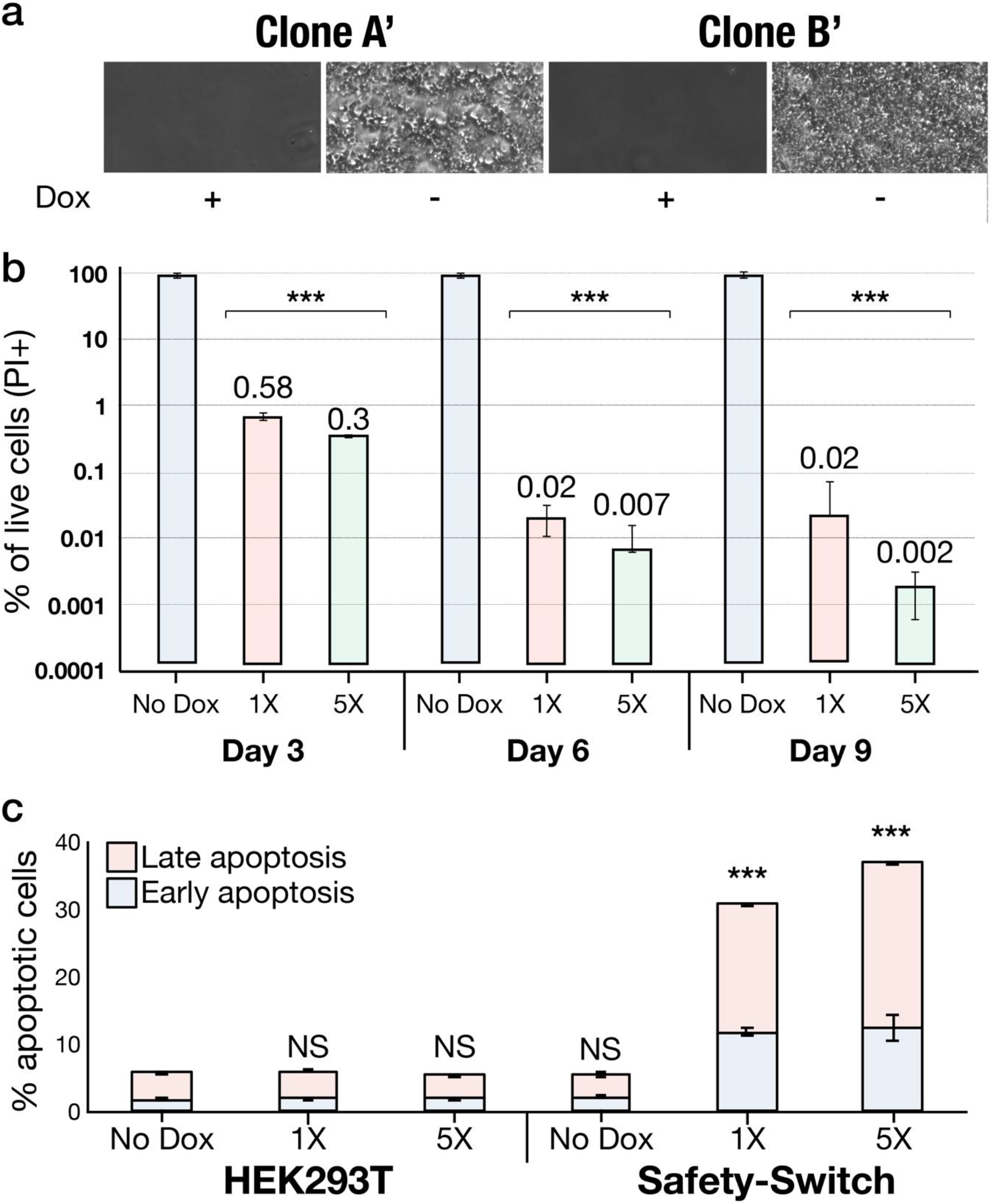
Towards the development of CRISPR Safety Switch. (**a**) Biosafety switch engineered HEK 293T treated with or without DOX, 10X microscope images of the cell lines A’ and B’ after 10 days of treatment. (**b**) Cell elimination sensitivity of the biosafety switch in HEK 293T: Percent of live cells (Propidium iodide positive) after 3, 6 or 9 days of treatment with 0, 1X or 5X of doxycycline (DOX). 1X of DOX corresponds to 1ug/mL and 5X to 5ug/mL. The Y-axis is in a logarithmic scale. (**c**) Evaluation of doxycycline toxicity and spontaneous activation of the biosafety switch in HEK 293T. Percent of cells in early or late apoptosis as assessed by propidium iodide and annexin V staining 3 days after treatment with 0, 1X or 5X of doxycycline (DOX). In all histograms, error bars represent standard error, n=3. Student t-tests were performed and marked NS, not significant (P > 0.05); ***P < 0.001 as compared to the positive controls in (b) and to the negative control in (c).

Under the microscope, both individual clones A’ and B’ showed almost complete clearance after 10 days of treatment with DOX. We next sought to make a quantitative analysis of the safety switch efficiency by counting the number of viable cells by flow cytometry with or without DOX treatment. To do so, we treated the clone B’ with 0 ug/mL, 1 ul/mL (1X) or 5 ul/mL (5X) of DOX and counted the number of live cells (propidium iodide positive cells) after 3, 6 or 9 days of treatments. After 3 days of treatment with 5X of DOX, the clonal population displayed a 99.7% reduction of cells as compared to the same population treated without DOX. The observed cell clearance went up to 99.98% after 9 days of treatment (Fig. 4b).

We have shown that the activation of our inducible circuit targeting repetitive elements is effective and results in the almost complete clearance of the cells. However, in order to consider implementing a safety switch in a clinical setting, the system, on top of being efficient would ideally have 1) little spontaneous action so that the transplanted cells stay viable and therefore keep their therapeutic benefits; and 2) the activating molecule should be inert and non-toxic. Therefore, we next investigated and quantified the self-activation or “leakiness” of our system when no DOX has been added and we assessed the cell toxicity triggered by DOX. We cultured and treated the B’ clonal population in addition to non-modified HEK 293T as a control with 0, 1X or 5X of DOX for 3 days and performed an annexin V – propidium iodide assay to quantify cell death in the different conditions (Fig. 4c). The control HEK 293T cells displayed basal apoptosis levels when treated with 1X or 5X of DOX, suggesting that both concentrations of the activating molecule are not toxic to the cells. Similarly, our safety switch engineered clone B’ treated with DOX did not display any abnormal apoptosis as compared to the control when 1X or 5X of DOX triggered respectively a 6- and a 7-fold increase in the apoptosis level as compared to the control cells.

Together these results suggest that 1) the safety switch is effective at eliminating the engineered cells to up to 99.98%, 2) DOX is not toxic to the cells, and 3) the system does not spontaneously activate itself in the absence of the activating molecule but only when it is added to the medium of culture.

## Discussion

In this study, we presented several biocontainment systems using lethal repetitive sgRNAs that induce rapid and robust cell death upon encountering a CRISPR/Cas9-based genome editor, but will be otherwise inert, as a prophylactic defense system against undesired genetic modification or as a stringent inducible safety-switch circuit.

Beyond the initial excitement and optimism surrounding CRISPR/Cas for its ease of use and positive applications lie the potential for dual use that demands awareness and motivates the development of protections and countermeasures. CRISPR-DS, the synthetic system we generated to prevent CRISPR-based genome editing ensures that introduction or activation of Cas9 triggers cell death, rendering cell populations in which the system is active effectively uneditable by Cas9. This system may either be activated in cells that have never been otherwise edited, rendering them safe from alteration, or after they have already gone through earlier rounds of editing, establishing “tamper-proof” information storage within a biological system. This biocontainment system would prevent edits to populations of cells by removing those that encounter Cas9, preventing them from passing on their genetic modification. As an alternative, we have shown that protein based anti-CRISPR approaches efficiently reduce Cas9 activity although they are still leaky and on population scales will be evaded by mutational escape while CRISPR-DS provides more stringent and persistent protection in such cases where prevention of undesired DNA editing is paramount.

To further enhance the utility of CRISPR-DS, the design of the lethal sgRNAs would ideally account for a wide range of potential genome editing effectors. In the case of known CRISPR/Cas9-based systems, sgRNA scaffolds specific to these systems are all that is required to protect against each category of enzyme. To adapt our genome editing prevention system to additional orthologues beyond SpCas9^24^, new sgRNA targets with compatible PAMs could be designed for *Staphylococcus aureus* Cas9, Cas12a or future Cas variants rapidly in response to their release (Suppl. Table 1). Utilizing evolutionarily conserved repetitive elements, a broad set of species may be covered by a relatively small number of sgRNA targets, while multiple orthologs of CRISPR/Cas9 may be included in the CRISPR Defense System to keep pace with the continuously expanding toolbox for genome editing.

As the use of Cas9 technologies for gene therapy is becoming more common, many therapeutic applications involve the use of *ex vivo* delivery of Cas9 to disrupt a target allele^25^ or precise correction by homologous recombination^26^. Repetitive sgRNAs may be used after an initial round of modification to negatively select cells that still contain gene editing reagents and could still potentially had the opportunity to generate undesired off-target mutations and secondary effects downstream.

For potential clinical applications, the CRISPR Safety-Switch could be leveraged as a highly escape-resistant biocontainment switch that would be activated if the host experiences complications such as host *vs* graft disease, cytokine storm, cancer or other unanticipated reaction from the modified cells. This bio-safety switch has shown to be highly effective at eliminating the cells in which the system is activated (up to 99.98% of the cells) with undetectable spontaneous activity and activated by a non-toxic molecule. In addition, our technology seems more stringent *in vitro* as compared to other available safety-switches. In 2012, a study carried out by Marin et al.^27^ benchmarked the following technologies: the Herpes simplex virus thymidine kinase (*HSV-TK*), the human inducible caspase 9 (*iCasp9*), the human *CD20* monoclonal antibodies and the mutant human thymidylate kinase (*mTMPK*). They assessed *in vitro* the efficiency of each technology, and observed 4 days after induction that the mean percent cell death was respectively, 94%, 94%, 96% and 70%. However, since our system includes Cas9, an exogenous protein coming from *S. pyogenes*, and even though its expression is repressed when the switch is OFF, we cannot exclude the potential immunogenicity of the biosafety switch in human.

We have shown that the genotoxicity driven by targeting repetitive elements of the genome can be leveraged to build mammalian synthetic biocontainment circuits such as CRISPR-DS and CRISPR Safety-Switch. As synthetic biology progresses towards grand challenges such as de-extinction and whole genome recoding proper defense against undesired genome edits or proper biocontainment become key ethical issues that must be addressed.

## Acknowledgements

We would like to thank John Aach, Chun-Ting Wu, Benedikt Markus and Mehdi Jafari for their useful advice throughout the research project.

## Funding

Research reported in this publication was supported by National Human Genome Research Institute of the National Institutes of Health under award number RM1HG008525 and by the Boehringer Ingelheim Fonds.

## Authors contribution

O.C., C.S., S.W., D.T, H.M., and G.M.C. initiated the study; O.C., C.S. designed the experiments; O.C., C.S., K.S., P.K., R.Y., and M.D. performed the experiments; O.C., C.S., P.K. and R.F. analyzed the data; O.C. and C.S. wrote the paper with contributing input from R.F., K.S., P.K., H.M. and G.M.C.

## Conflict of interests

G.M.C. is a co-founder of Editas Medicine and has other financial interests listed at arep.med.harvard.edu/gmc/tech.html.

## Data and materials availability

All NGS data used for the main and supplementary figures have been made available at SRA BioProject Accession PRJNA604267 for Fig. 1B, PRJNA604277 for Fig. 3A and PRJNA604281 for Fig. S1.

## References

1. Cui, L. & Bikard, D. Consequences of Cas9 cleavage in the chromosome of Escherichia coli. Nucleic Acids Res. 44, 4243–4251 (2016).

2. Reisch, C. R. & Prather, K. L. J. The no-SCAR (Scarless Cas9 Assisted Recombineering) system for genome editing in Escherichia coli. Sci. Rep. 5, (2015).

3. Citorik, R. J., Mimee, M. & Lu, T. K. Sequence-specific antimicrobials using efficiently delivered RNA-guided nucleases. Nat. Biotechnol. 32, 1141–1145 (2014).

4. Ceccaldi, R., Rondinelli, B. & D’Andrea, A. D. Repair Pathway Choices and Consequences at the Double-Strand Break. Trends Cell Biol. 26, 52–64 (2016).

5. Mladenov, E., Magin, S., Soni, A. & Iliakis, G. DNA double-strand-break repair in higher eukaryotes and its role in genomic instability and cancer: Cell cycle and proliferation-dependent regulation. Semin. Cancer Biol. 37–38, 51–64 (2016).

6. Roos, W. P. & Kaina, B. DNA damage-induced cell death by apoptosis. Trends Mol. Med. 12, 440–450 (2006).

7. Aguirre, A. J. et al. Genomic Copy Number Dictates a Gene-Independent Cell Response to CRISPR/Cas9 Targeting. Cancer Discov. 6, 914–929 (2016).

8. Kuscu, C. et al. CRISPR-STOP: gene silencing through base-editing-induced nonsense mutations. Nat. Methods (2017) doi:10.1038/nmeth.4327.

9. Yang, L. et al. Genome-wide inactivation of porcine endogenous retroviruses (PERVs). Science 350, 1101–1104 (2015).

10. Xing, J., Witherspoon, D. J. & Jorde, L. B. Mobile element biology: new possibilities with high-throughput sequencing. Trends Genet. 29, 280–289 (2013).

11. Venter, J. C. et al. The Sequence of the Human Genome. Science 291, 1304–1351 (2001).

12. Lander, E. S. et al. Initial sequencing and analysis of the human genome. Nature 409, 860–921 (2001).

13. de Koning, A. P. J., Gu, W., Castoe, T. A., Batzer, M. A. & Pollock, D. D. Repetitive Elements May Comprise Over Two-Thirds of the Human Genome. PLoS Genet. 7, (2011).

14. Smith, C. J. et al. Enabling large-scale genome editing by reducing DNA nicking. bioRxiv 574020 (2019) doi:10.1101/574020.

15. Penzkofer, T. et al. L1Base 2: more retrotransposition-active LINE-1s, more mammalian genomes. Nucleic Acids Res. 45, D68–D73 (2017).

16. Batzer, M. A. & Deininger, P. L. Alu repeats and human genomic diversity. Nat. Rev. Genet. 3, 370–379 (2002).

17. Rauch, B. J. et al. Inhibition of CRISPR-Cas9 with Bacteriophage Proteins. Cell 168, 150–158.e10 (2017).

18. Daley, G. Q. The Promise and Perils of Stem Cell Therapeutics. Cell Stem Cell 10, 740–749 (2012).

19. Chavez, A. et al. Highly efficient Cas9-mediated transcriptional programming. Nat. Methods 12, 326–328 (2015).

20. Brinkman, E. K., Chen, T., Amendola, M. & van Steensel, B. Easy quantitative assessment of genome editing by sequence trace decomposition. Nucleic Acids Res. 42, e168 (2014).

21. Byrne, S. M. & Church, G. M. Crispr-mediated Gene Targeting of Human Induced Pluripotent Stem Cells. Curr. Protoc. Stem Cell Biol. 35, 5A.8.1–22 (2015).

22. Park, J., Lim, K., Kim, J.-S. & Bae, S. Cas-analyzer: an online tool for assessing genome editing results using NGS data. Bioinformatics 33, 286–288 (2017).

23. Ferreira, R., Gatto, F. & Nielsen, J. Exploiting off-targeting in guide-RNAs for CRISPR systems for simultaneous editing of multiple genes. FEBS Lett. 591, 3288–3295 (2017).

24. Cebrian-Serrano, A. & Davies, B. CRISPR-Cas orthologues and variants: optimizing the repertoire, specificity and delivery of genome engineering tools. Mamm. Genome 28, 247–261 (2017).

25. Zhang, Y. et al. CRISPR-Cas9 mediated LAG-3 disruption in CAR-T cells. Front. Med. (2017) doi:10.1007/s11684-017-0543-6.

26. Zhu, P. et al. CRISPR/Cas9-Mediated Genome Editing Corrects Dystrophin Mutation in Skeletal Muscle Stem Cells in a Mouse Model of Muscle Dystrophy. Mol. Ther. Nucleic Acids 7, 31–41 (2017).

27. Marin, V. et al. Comparison of Different Suicide-Gene Strategies for the Safety Improvement of Genetically Manipulated T Cells. Hum. Gene Ther. Methods 23, 376–386 (2012).

