## Supplementary Figures for "CRISPR-mediated biocontainment"

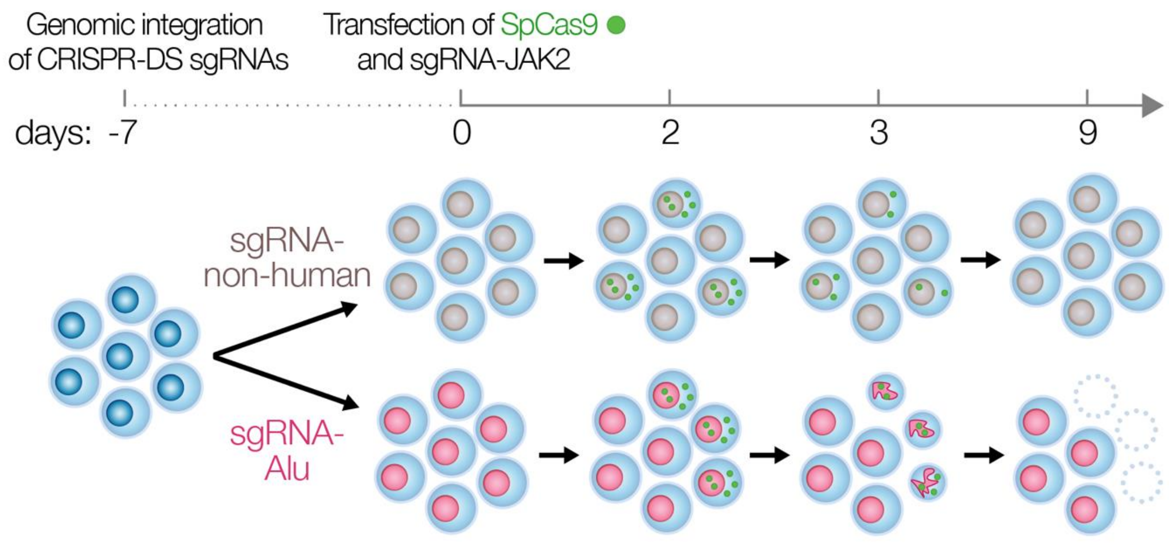


**Figure S1 | Visual depiction of the genomic integration of the CRISPR-DS system and experimental overview.** Active CRISPR-DS expressing sgRNA-Alu is shown in red while the control (non-functional) CRISPR-DS is shown in light brown expressing sgRNA-non-human. Annexin V+ Propidium iodide+ FACS populations.

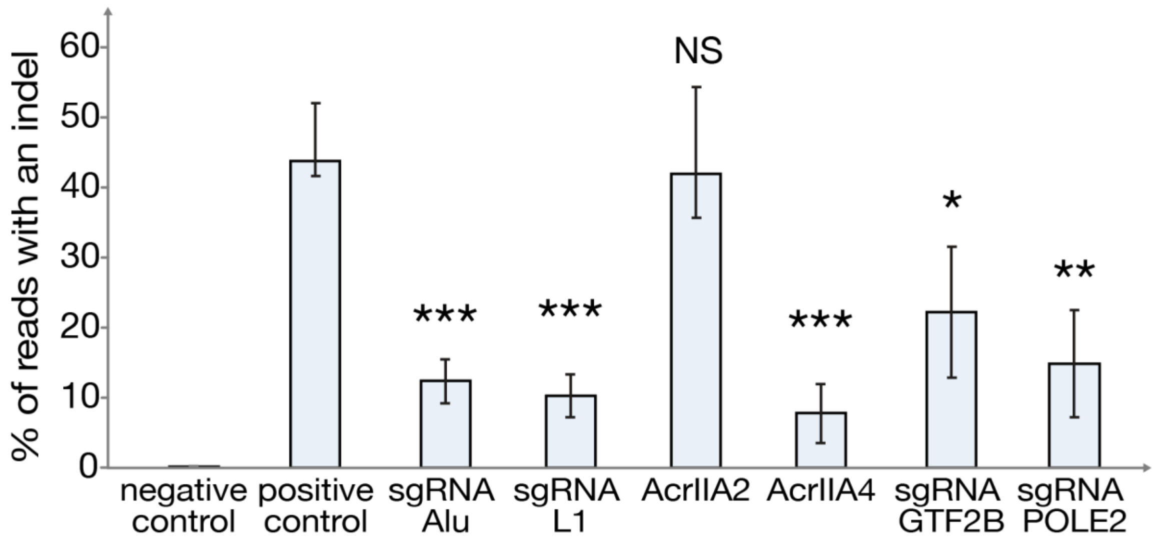


**Figure S2 | CRISPR-DS compared to systems targeting essential genes or using anti-CRISPR proteins.** Prevention of DNA modification at the JAK2 locus by CRISPR-DS in HEK 293T cell line transfected with either repetitive element sgRNA, anti-CRISPR plasmids, or an essential gene sgRNA. The graph represents the mean of three biological replicates for indel mutation rate at the JAK2 locus three days after transfection, which is plotted on the y-axis. In all histograms, error bars represent standard error, n=3. Student t-tests were performed and marked NS, not significant ($P > 0.05$); *$P < 0.05$; **$P < 0.01$; ***$P < 0.001$ as compared to the positive control.
